# Prediction of pyrazinamide resistance in *Mycobacterium tuberculosis* using structure-based machine learning approaches

**DOI:** 10.1101/518142

**Authors:** Joshua J Carter, Timothy M Walker, A Sarah Walker, Michael G. Whitfield, Glenn P. Morlock, Charlotte I. Lynch, Dylan Adlard, Timothy EA Peto, James E. Posey, Derrick W Crook, Philip W Fowler

## Abstract

**Background:** Pyrazinamide is one of four first-line antibiotics used to treat tuberculosis, however antibiotic susceptibility testing for pyrazinamide is challenging. Resistance to pyrazinamide is primarily driven by genetic variation in *pncA,* an enzyme that converts pyrazinamide into its active form.

**Methods:** We curated a dataset of 664 non-redundant, missense amino acid mutations in *pncA* with associated high-confidence phenotypes from published studies and then trained three different machine learning models to predict pyrazinamide resistance. All models had access to a range of protein structural-, chemical- and sequence-based features.

**Results:** The best model, a gradient-boosted decision tree, achieved a sensitivity of 80.2% and a specificity of 76.9% on the hold-out Test dataset. The clinical performance of the models was then estimated by predicting the binary pyrazinamide resistance phenotype of 4,027 samples harboring 367 unique missense mutations in *pncA* derived from 24,231 clinical isolates.

**Conclusions:** This work demonstrates how machine learning can enhance the sensitivity/specificity of pyrazinamide resistance prediction in genetics-based clinical microbiology workflows, highlights novel mutations for future biochemical investigation, and is a proof of concept for using this approach in other drugs.

## Introduction

*Mycobacterium tuberculosis* is an evolutionarily ancient human pathogen that is the leading cause of death by infectious disease worldwide, except during the SARS-CoV-2 pandemic. In 2021, tuberculosis was responsible for 1.6 million deaths and 10.6 million new infections^1^. Tuberculosis control efforts have been hampered by the evolution of resistance to antibiotics, threatening the efficacy of the standard four drug antibiotic regimen consisting of rifampicin, isoniazid, ethambutol, and pyrazinamide. Pyrazinamide plays a critical role in tuberculosis treatment through its specific action on slow-growing, “persister” bacteria that often tolerate other drugs due to their reduced metabolism^2–6^. Due to its unique sterilizing effect and its synergy with new tuberculosis drugs such as bedaquiline, pyrazinamide is also included in new treatment regimens targeting drug-resistant tuberculosis^7–12^. Therefore, accurately and rapidly determining whether a clinical isolate is resistant to pyrazinamide is critically important for the treatment of tuberculosis.

Most culture-based laboratory methods to determine pyrazinamide resistance are technically challenging, requiring highly trained technicians. Even then, results are often not reproducible, meaning these methods are rarely employed in low-resource and/or high-burden clinical settings^13^. Even the current gold standard, the Mycobacteria Growth Indicator Tube (MGIT), which is relatively simple to use, can suffer from low precision, with false-resistance rates of 1-68% reported^14–20^. As the prevalence of multidrug-resistant and extensively drug-resistant TB increases, this lack of precision will become more of a problem.

Resistance to rifampicin or isoniazid can be predicted in most isolates (90-95% and 50-97%, respectively) by the presence of a small number of highly-penetrant genetic variants in short and well-delineated regions of one or two genes (*rpoB* and *katG*/*fabG1*, respectively)^3^. However, despite pyrazinamide being used to treat tuberculosis since 1952, comparatively less is known about which genetic variants confer resistance compared to other first-line drugs^4^. In the recent catalogue of resistance-associated mutations of *M. tuberculosis* published by the World Health Organization, the performance for pyrazinamide was markedly lower (72.3% sensitivity and 98.8% specificity) than either rifampicin or isoniazid (93.8% and 98.2% or 91.2% and 98.4% sensitivity and specificity respectively)^21,22^. While some of this poor performance is likely due to inaccuracies in phenotypic testing, a comprehensive genetic catalogue for pyrazinamide resistance mutations remains elusive.

Pyrazinamide is a pro-drug that is converted to its active form of pyrazinoic acid by the action of PncA, a pyrazinamidase/nicotinamidase encoded by the *pncA* gene^23^. While other genetic loci have been implicated in pyrazinamide resistance (notably *rpsA*, *panD*, *clpC1*, and the putative efflux pumps *Rv0191*, *Rv3756c*, *Rv3008*, and *Rv1667c*), the majority (70-97%) of pyrazinamide-resistant clinical isolates harbor genetic variants in either the promoter region or coding sequence of *pncA*^21,24–34^. In contrast to the well-delineated and relatively restricted “resistance-determining regions” found in *rpoB* (rifampicin, 27 codons) and *katG* (isoniazid, single codon), pyrazinamide-resistant variants have been identified along the entire length of the *pncA* gene (**Figure 1A**) with no single variant predominating. Hence, while targeted- or whole-genome sequencing approaches are capable of assaying the entire *pncA* gene, the number and diversity of resistance-conferring variants in *pncA* fundamentally limits the sensitivity and specificity of heuristic approaches that aim to predict the effectiveness of pyrazinamide based on a catalogue of previously-observed genetic variants^3,13,24,29,30,35,36^.

**Figure 1.**
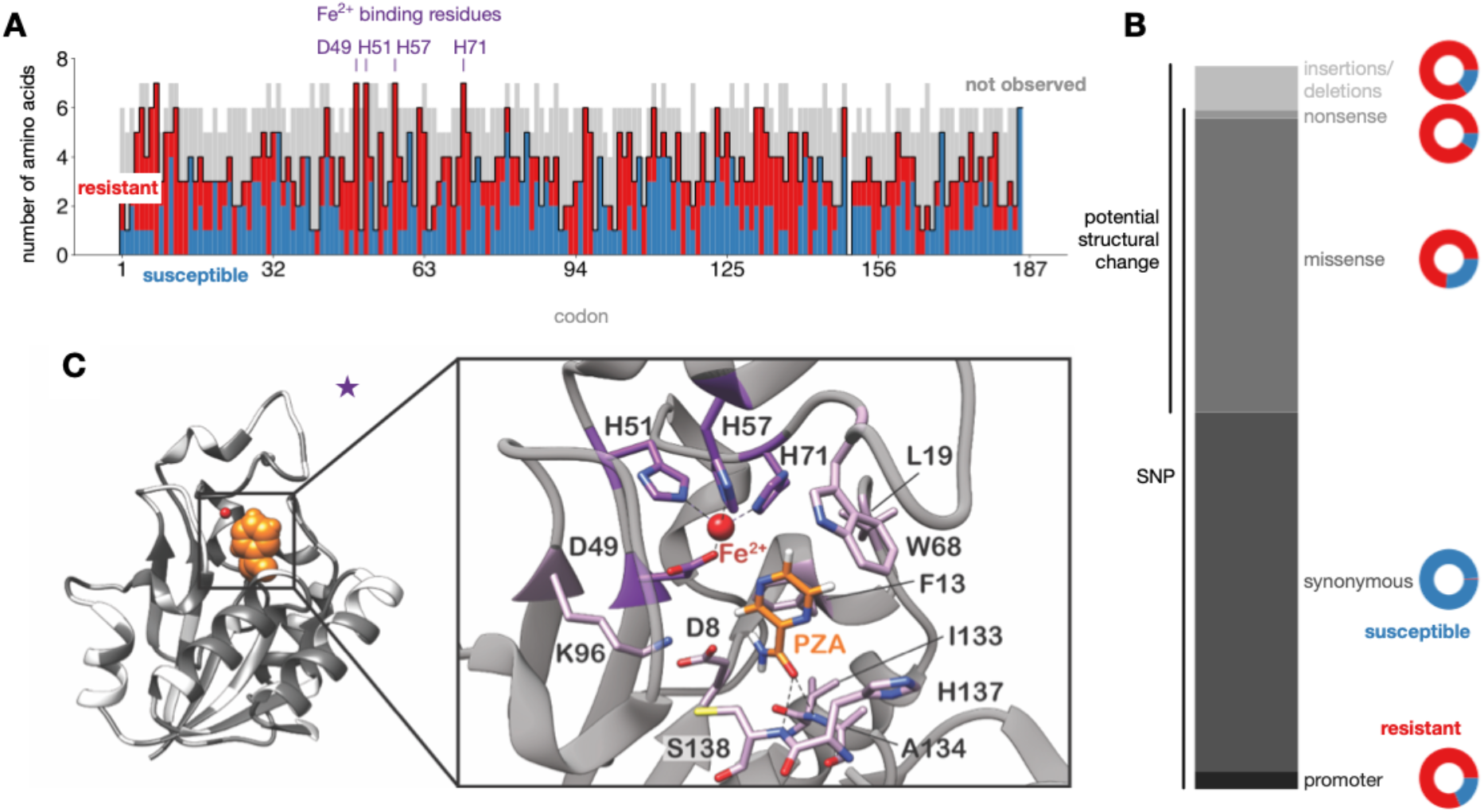
Distribution of PncA mutations from published datasets. (**A**) Barplot of the impact of possible missense mutations in PncA by amino acid position. High confidence resistant (red) and susceptible (blue) mutations are overlaid on the possible missense mutations whose effect on resistance is unknown or unclear (grey). (**B**) Distribution of the types of mutations reported by the CRyPTIC consortium *et al*. (**C**) Missense mutations from the dataset plotted onto the PncA structure (PDB ID: 3PL1) in dark grey. A pyrazinamide molecule (orange) has been modeled into the active site.

Genetics-based clinical microbiology for tuberculosis currently depends on being able to *infer* the effect of any likely occurring *pncA* mutation on pyrazinamide susceptibility. Recent studies to identify pyrazinamide-resistance determining mutations have focused on either classifying mutations from previously observed clinical isolates or discovering novel mutations through *in vitro/in vivo* screening approaches^21,22,30,37–39^. However, these strategies are constrained, respectively, by the relative paucity of sequenced clinical isolates compared to the number of potential resistance-causing mutations and the lack of laboratory capacity to systematically generate and test mutants. Computational modelling approaches^40^ can potentially *predict* the effect of a significant number of missense mutations^41–44^ before they are observed in clinical isolates. Several studies have already trained machine learning models on a number of anti-tuberculars^45–48^, including pyrazinamide^49^.

As PncA is not essential and can be inactivated through defects in protein folding, reduced stability, distortion of active site geometry, abrogation of metal binding, or some combination of these, we expected a machine-learning approach to be ideally suited to simultaneously consider all these possible mechanisms of PncA inactivation, and hence more accurately predict pyrazinamide resistance/susceptibility. In this paper, we confirm using the largest *Train/Test* and *Validation* datasets used to date that machine-learning models that learn from a range of structural, chemical and evolutionary features can robustly and accurately predict the effect of missense amino acid mutations on pyrazinamide susceptibility.

## Materials and Methods

We first constructed independent *Train, Test* and *Validation* datasets (**Table 1**). The first two were built by combining a comprehensive *in vitro*/*in vivo* mutagenesis study^38^ with two published catalogues of *M. tuberculosis* genetic variants associated with resistance^21,39^ resulting in a *Train/Test* dataset of 664 non-redundant missense mutations (349 associated with resistance) where there was no discrepancy in the predicted phenotype. This was then split 70:30 to produce independent *Train* and *Test* datasets containing 464 and 200 mutations, respectively (**Methods, Table 1)**. The *Validation* dataset was constructed by aggregating 24,231 clinical samples from three collections^13,30,50^ resulting in 4,027 samples containing one of 367 non-redundant missense PncA mutations. Briefly, phenotypes for strains with single missense mutations in *pncA* were aggregated by mutation, tallying the results of the phenotypic testing. Mutations that were resistant or susceptible at least 75% of the time and that had been phenotyped at least 4 times were included. Additionally, mutations that had been phenotyped at least twice with no discrepancies were also added. Finally, a further independent *Quantitative*

**Table 1.** Description of datasets employed in this study. (R=resistant to antibiotic, S=susceptible, U=inconsistent results, MIC=Minimum inhibitory concentration) dataset was created by measuring the minimum inhibitory concentration (MIC) on a small number of missense mutations to test if our models can predict the magnitude of the effect.

| Dataset | Phenotype | # Isolates | # non-redundant missense mutations |
| --- | --- | --- | --- |
| *Train* | R/S | n/a | 464 |
| *Test* | R/S | n/a | 200 |
| *Validation* | R/U/S | 24,231 | 367<br>(199 with an R/S phenotype) |
| *Quantitative* | MIC | 71 | 57 |

### Pyrazinamide minimum inhibitory concentration determination

Isolates used for MIC determination came from the EXIT-RIF study and US Centers for Disease Control. Of the 366 *Mycobacterium tuberculosis* clinical isolates, 333 were collected as part of a prospective cohort study (“EXIT-RIF”) between November 2012 and December 2013 in three South African provinces (Free State, Eastern Cape and Gauteng). A *Mycobacterium tuberculosis* databank housed at the SAMRC Centre for Tuberculosis Research, consisting of ∼45,000 drug resistant isolates collected in the Western Cape province since 2001, was queried to identify isolates containing both PZA MIC data and *pncA* genotypic data, this produced the remaining 33 *Mycobacterium tuberculosis* clinical isolates. Isolates that harbored single amino acid substitutions in PncA (39 out of 366 total) were selected for comparison to model predictions. An additional 32 clinical isolates (collected from 2000 to 2008) harboring single missense mutations in *pncA* came from the culture collection at the Laboratory Branch, Division of Tuberculosis Elimination, US CDC.

All MICs were determined using the non-radiometric BACTEC MGIT 960 method (BD Diagnostic Systems, NJ, USA) with manufactured-supplied pyrazinamide medium/supplement as previously described^51^. This system makes use of modified test media which supports the growth of mycobacteria at a pH of 5.9. Isolates from the EXIT-RIF study were tested at 100, 75, 50, 25 µg/ml whilst the Center for Disease Control used PZA concentrations of 50, 100, 200, 300, 400, 600 and 800 µg/ml. A fully susceptible MTB laboratory strain H37Rv (ATCC 27294) was included as a control for all isolates tested. The resulting 71 isolates contained one of 59 missense mutations; of the ten mutations measured more than once, two had inconsistent phenotypes and were removed, leaving 57 missense mutations of which 50 were resistant and 7 susceptible.

### Determination of structural-, chemical- and evolutionary features

Since the *M. tuberculosis* structure of PncA^52^ (PDB: 3PL1) does not contain electron density for pyrazinamide, we first fitted it onto the *A. baunmanii* PncA structure^53^ and retained the coordinates of PZA. A wide range of structural, chemical, thermodynamic and evolutionary features were added^54^, including the change in mass, volume, isoelectric point, hydrophobicity and chemistry^55^ and the distances from the Fe^2+^ ion and pyrazinamide molecule, solvent accessibility, backbone angles, secondary structure, temperature factor, depth and degree of hydrogen bonding. To assess a *pncA* mutation’s impact on the stability of PncA, we added scores from three meta-predictors (RaSP^56^, mCSM^57^ and DeepDDG^58^). Finally we added MAPP scores, which aims to quantify the evolutionary constraints imposed on a given position in a protein^59^, and SNAP2 scores. SNAP2 is a neural network trained to predict whether protein mutations are neutral or have a deleterious effect on function^60^.

### Training and Reproducibility

Logistic regression (LR), a multi-layer perceptron classifier (NN) and a gradient-boosted decision tree (XB) were all trained as described in our online code and data repository – this also contains saved states of the final models and Python3 jupyter-notebooks allowing one to reproduce in a web browser all results and figures^61^.

## Results

### Observed genetic variation in pncA

Since it includes the results of an *in vitro* mutagenesis study, the *Train/Test* dataset captures the most genetic variation in *pncA*. Mutations are observed at every codon bar one (**Figure 1A**) and all possible amino acids arising from a single nucleotide substitution are observed at several codons. Interestingly, there were a significant number of *pncA* codons where mutations associated with either resistance or susceptibility were seen, confirming that the change in local chemistry introduced by the mutant amino acid is an important factor in determining resistance (**Figure 1A**). The codons with the greatest mutational diversity in the dataset were all residues involved in active site formation or metal binding, suggesting that, consistent with our hypothesis, loss or alteration of these functions is a common mechanism for gaining pyrazinamide resistance. Indeed, previous studies have noted a negative correlation between a mutation’s distance from the active site and its tendency to cause resistance^30,38,52^.

### Clinically-observed association between genetic variation in pncA and pyrazinamide resistance

Overall 3,351 samples (14.7%) in the CRyPTIC dataset are resistant to pyrazinamide and 6,851 samples have one or more genetic variants in either the promoter and/or open reading frame of *pncA*. The majority (6,622 samples / 96.7%) have a single genetic variant with 93.9% (6,221 samples) of these being substitutions. The remaining 401 samples (6.1%) contained insertions, deletions and frameshifts and these were strongly associated with resistance (343 samples, 85.5%)^21,39^, consistent with their likely disruption of the PncA enzyme. Most synonymous substitutions (present in 3,288, 49.7% of the single variant strains, **Figure 1B**) were not associated with resistance, however seven variants were observed in resistant isolates. S65S (19 resistant isolates) is a phylogenetic SNP present in Lineage 1; however, it is susceptible in 3,204 strains, suggesting that these 19 isolates are either phenotyping errors or that there is an alternative mechanism of pyrazinamide resistance at play in these strains. The remaining mutations—R2R, L19L, A46A, D63D, 131V and V155V—are each present a single time (twice for A46A), limiting our ability to associate these variants with resistance. Thus, non-synonymous substitution variants (present in 2,766, 41.8% of single variant strains) appear to be associated with most of the pyrazinamide resistance in *M. tuberculosis*.

### Feature determination using Test/Train dataset

To understand the structural features that determine a mutation’s effect on pyrazinamide susceptibility, we mapped our combined *Train/Test* dataset onto the PncA structure. No obvious clustering was revealed, consistent with the previously observed distribution of resistant mutations across the gene sequence and protein structure (**Figure 1A,C**) ^13,24,29,30,38^. Examining the PncA structure also suggested that resistant mutations were more likely to be buried in the hydrophobic core of the protein and therefore likely destablising, consistent with findings from previous *in vitro* and *in vivo* screens (**Figure 2A**)^30,38^. Indeed, some pyrazinamide-resistant mutations result in reduced production of functional PncA, perhaps due to impaired protein folding/stability^38,62^. Despite having a similar learning objective there was only a moderate level of correlation between the different models that predicted the effect of a mutation on the protein stability (**Figure 2B**). Other more accurate methods exist, but these require several orders of magnitude of computational resource^63^. Since SNAP2 uses evolutionary information derived from a multiple sequence alignment, one might expect some similarity to MAPP, but again there is only a moderate degree of correlation between the two scores (**Figure 2B**).

**Figure 2:**
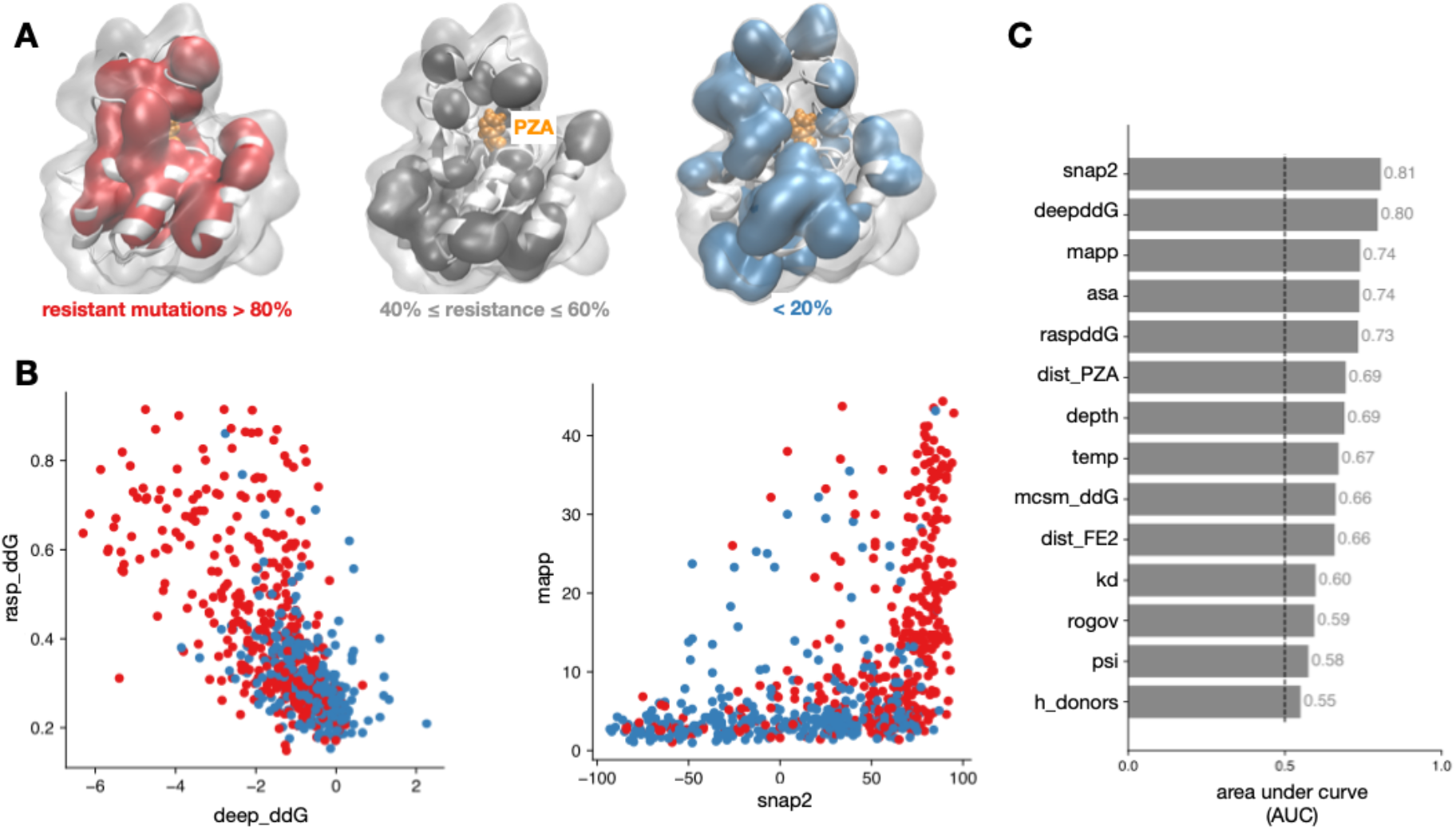
Structural and evolutionary traits correlate with mutational impact on pyrazinamide susceptibility. (**A**) Amino acids where >80% of mutations confer resistance are more likely to be found in the core of PncA. (**B**) There is only a moderate correlation between RaSP & DeepDDG, which both predict the effect of a mutation on protein stability, and MAPP and SNAP2. Resistant and susceptible mutations are plotted in red and blue, respectively. (**C**) The performance of individual features, as measured by the area under curve (AUC) of the receiver operator characteristic of a univariable logistic regression. The dashed line denotes random guessing.

### Machine-learning models accurately predict pyrazinamide resistance

Univariable logistic regression over the derivation dataset revealed that most of the individual predictors were associated with resistance (**Figure 2C, S1**). The SNAP2 score and DeepDDG protein stability scores proved to be the most discriminatory individual features and six features (change in molecular weight, volume and isoelectric point along with the secondary structure, φ backbone angle and number of hydrogen bond acceptors) were discarded at this point since their AUC lay below an arbitrary threshold of 0.55.

Following hyperparameter tuning, three different machine learning models (logistic regression, LR, a gradient-boosted decision tree, XB and a single layer neural network, NN) were trained on the *Train* dataset using 10-fold cross-validation (**Methods**). All three models performed similarly when applied to the *Train* dataset (**Figure 3**) with sensitivities of 78-79% and specificities in the range 83-86%. As expected, the models performed less well on the *Test* dataset and the LR model had a superior sensitivity (81.1%) than both the XB (80.2%) and the LR (73.8%) model, whilst the XB model had an improved specificity (76.9%) than either the LR or NN models (69.9% & 56.5%, respectively). We conclude that the gradient-boosted decision tree (XB) model performed best since it resulted in the fewest number of resistant samples incorrectly classified as susceptible (so-called very major errors, VMEs) and had the highest diagnostic odds ratio (**Figure 3B,C**).

**Figure 3:**
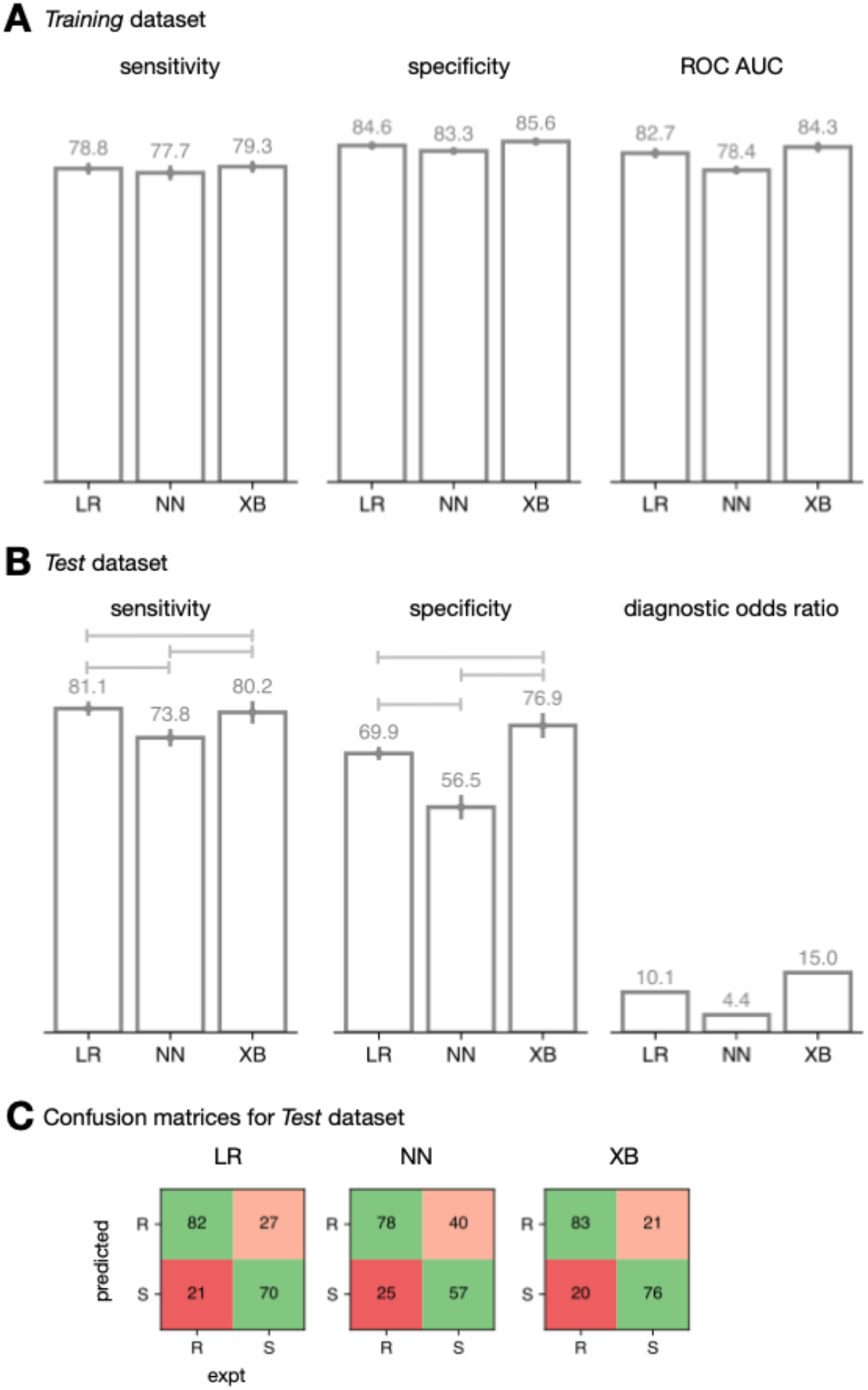
Machine learning models predict pyrazinamide resistance from structural, chemical and evolutionary features. Performance of logistic regression (LR), a simple neural network (NN) and gradient-boosted decision tree (XB) models on the (**A**) *Training* and (**B**) *Test* sets. Error bars represent 95% confidence intervals from bootstrapping (n=10) and brackets indicate a significant difference (z-test, *p* < 0.05) (**C**) Confusion matrices are shown for the *Test* set. Very major errors (VME, predicted S but R) are considered worse than major errors (ME, predicted R but S) and hence VMEs and MEs are shaded red and pink, respectively.

### Most residues that were incorrectly predicted as susceptible are surface-exposed

The models predicted 20-25 VMEs and misclassified a further 21-40 susceptible samples as resistant (major errors, ME). Collectively 12 VMEs and 11 MEs were shared between all three models (**Figure 4A**). Although the mutations responsible for the shared VMEs were dispersed throughout the protein structure, most (11/12) were surface exposed (**Figure 4B**). All these mutations were predicted by DeepDDG, mCSM and RasP to minimally decrease the stability of PncA compared to mutations correctly predicted to confer resistance, suggesting these errors may be partly due to inaccuracies in the predicted free energy change of unfolding (**Figure 4C**), although other features also contributed (**Figure S2**). Major errors were also dispersed throughout the protein and were more likely to be buried and to be predicted by SNAP2 to not have a functional effect compared to mutations correctly predicted to have no effect (**Figure 4C**), although again other features played a part (**Figure S2**).

**Figure 4.**
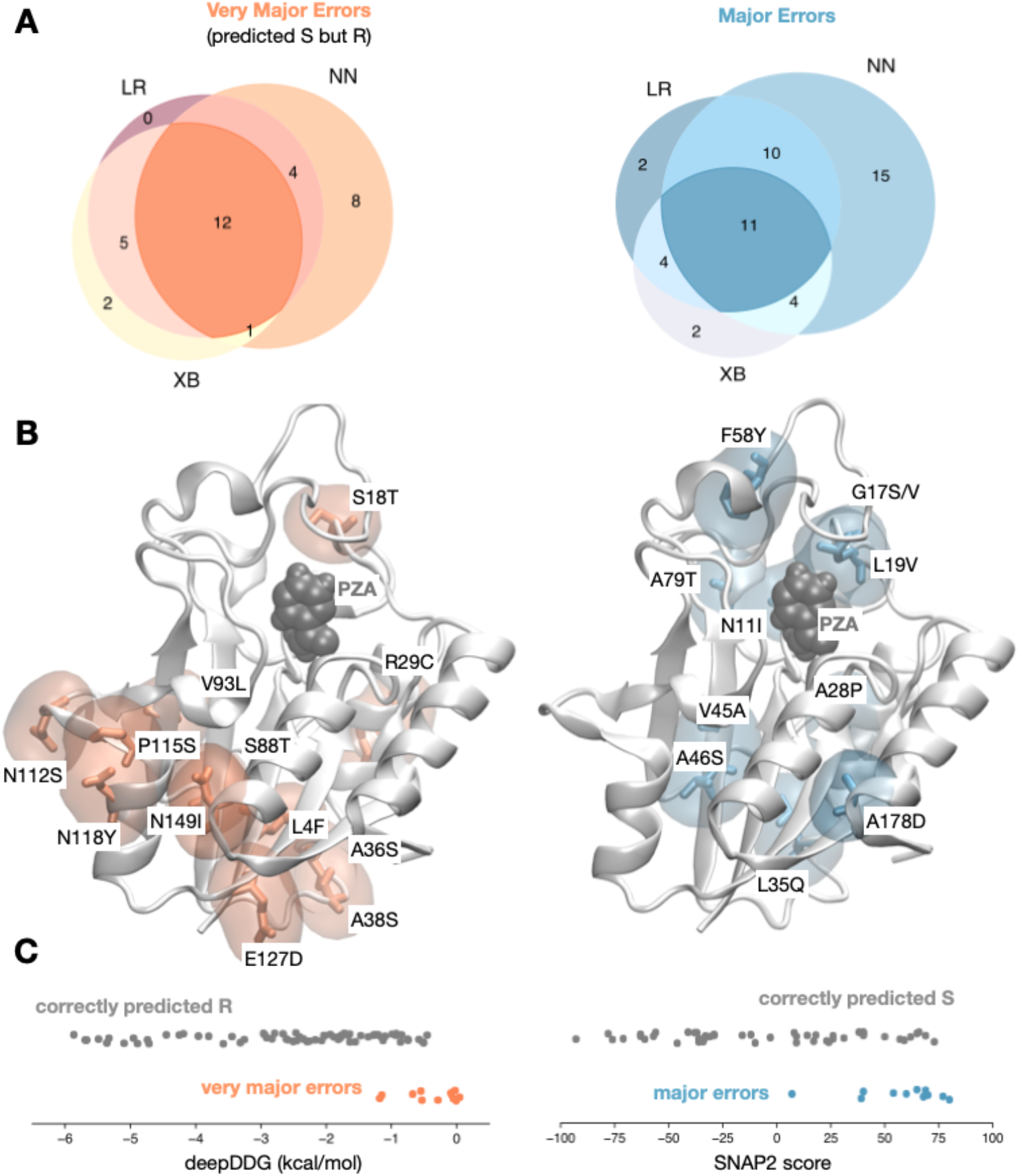
Very major errors are concentrated on the surface of PncA. (**A**) The majority of very major (VME) and major (ME) errors are shared between the three models. (**B**) PncA with the corresponding residues highlighted where the shared very major (orange) and major (blue) errors are found. (**C**) The shared very major and major errors are predicted to have less and more effect, respectively, on the stability of the protein, as exemplified by DeepDDG and the function of the protein, according to SNAP2.

Examining the feature importances of the gradient-boosted decision tree (XB) models (**Figure S3**) shows that whilst all 16 features are incorporated to some extent, the first four are all scores from other machine learning models (MAPP, DeepDDG, RaSP and SNAP2), with the next four all being derived from the protein structure (ψ backbone angle, residue depth and residue solvent accessible surface area) or describing the change in chemistry^55^.*Gradient-boosted decision tree model predictions generalize to a large clinical dataset*

A *Validation* dataset was derived from 24,231 *pncA* gene sequences with MGIT antibiotic susceptibility results (**Table 1**). Most samples contained no mutations in *pncA*: only 4,027 samples had one of 367 missense mutations. We assume this dataset is representative of the genetic diversity in PncA existing in clinical infections but it is likely biased due to oversampling of outbreak strains and other factors. Until very large unselected clinical datasets are collected and made publicly available, however, it is the best dataset available.

Applying the gradient-boosted decision tree (XB) model to this dataset (**Figure 5A**) resulted in a high sensitivity (97.2%) but a modest specificity (46.0%). The presence of a substantial number of samples in this dataset (908 samples, 22.5%) contained one of 168 (45.8%) mutations that either were only observed once, or whose phenotype varied between isolates was a key contributor to this reduction in performance. Whilst this dataset therefore captures the real-world variability of culture-based phenotypic methods for pyrazinamide susceptibility testing, it is not a good basis on which to assess performance and removing these samples improved the specificity to 63.1% (**Figure 5B**). Slightly over half (116, 58.3%) of remaining mutations were also present in the *Train* dataset and accounted for 2,044 out of the remaining 3,119 samples. The predictions for the samples in this group had a sensitivity of 98.3% and a specificity of 75.6%. As expected, the other 83 mutations found in 1,075 samples had a lower performance, with the specificity notably being 22.9%. Examining the performance at the level of the mutations (rather than samples) yields a specificity of 48.0%, however the size of the dataset is now small with only 25 out of 83 mutations having a susceptible phenotype. The XB model also outperforms a previously published model^49^ applied to this same dataset; SUSPECT-PZA achieved a sensitivity of 93.7% and a specificity of 44.3% on the original 4,027 samples. Only considering the 199 mutations with a consistent phenotype improved the specificity to 47.7% with a slight fall in sensitivity (92.3%), however this is less predictive than the performance of the gradient boosted decision tree.

**Figure 5.**
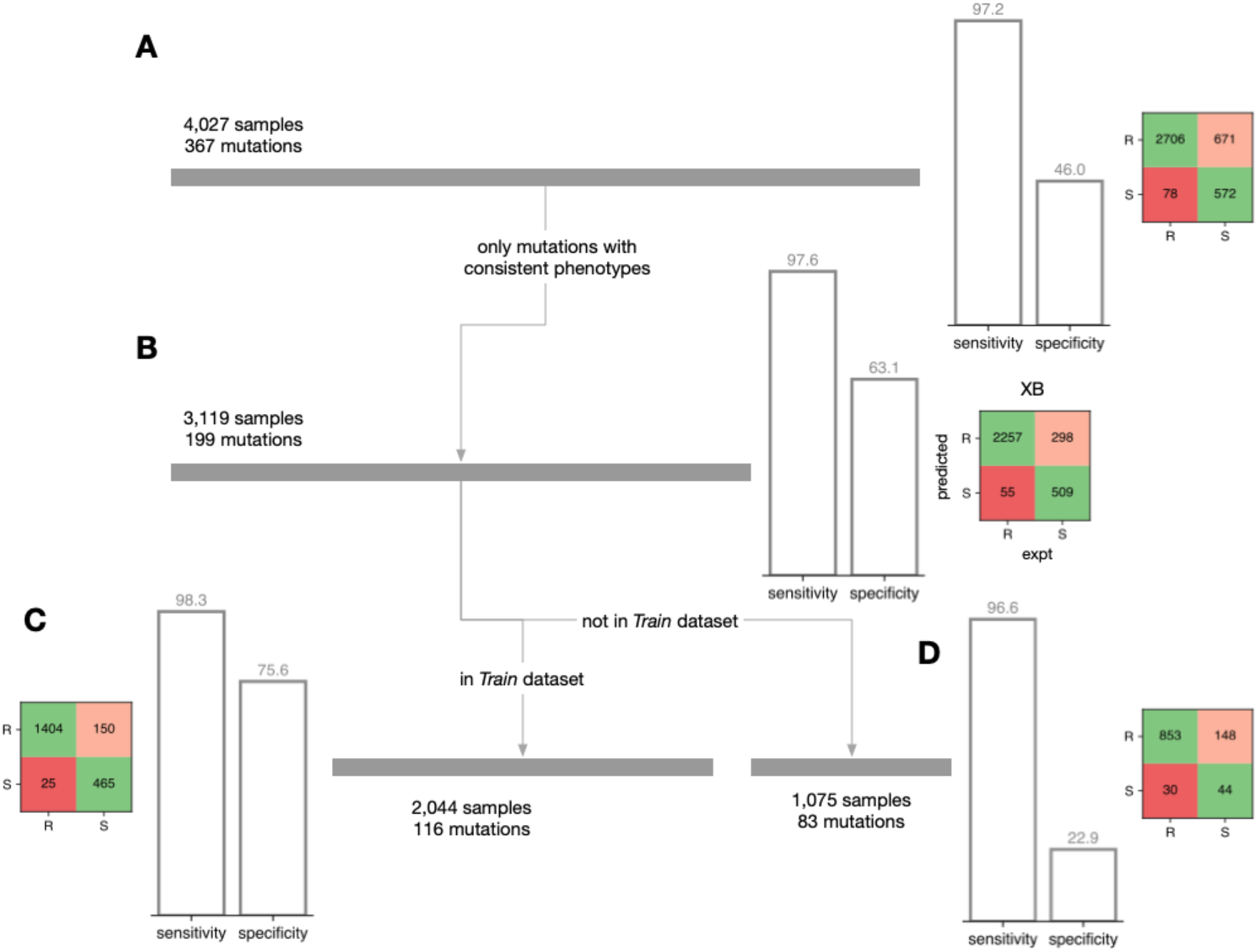
Performance on a real set of clinical samples. (**A**) Whilst the sensitivity is high, the specificity of the gradient-boosted decision tree model on the *Validation* dataset is lower than observed on the *Test* dataset. (**B**) Removing samples containing a mutation that has an experimentally inconsistent phenotype increases the specificity. As expected, splitting into samples whose mutation either (**C**) belongs or (**D**) does not belong to the Train dataset further stratifies performance.

### Comparison of model predictions with pyrazinamide minimum inhibitory concentrations in vitro

Since it is difficult to assess how much of the discordance in the previous section can be attributed to either error in the measured clinical phenotype or deficiencies in our model, we compared its predictions to minimum inhibitory concentration (MIC) data taken from a small but high-quality dataset of 71 *M. tuberculosis* isolates (59 unique missense mutations, quantitative dataset, **Methods**). This also enabled us to test the model’s capacity to predict the *degree* of pyrazinamide resistance conferred by a particular mutation, by comparing the calls and predicted probabilities of our model with the pyrazinamide MICs. Overall, our model correctly predicted the binary (R/S) phenotype for 51 of 57 missense mutations in PncA (**Figure S4**), and, crucially, predicted the correct phenotype for 6 out of 8 mutations that were not in either the *Train* or *Test* datasets. Utimately, many more samples with a wide range of pyrazinamide MICs will be needed to accurately assess if quantitative prediction is possible for this drug.

## Discussion

We have shown that machine learning models trained on structural, chemical and evolutionary features can predict whether missense amino acid mutations in *pncA* confer resistance to pyrazinamide, adding to the growing body of work that is exploring different ways of *predicting* antibiotic resistance from genetics^41–49^. While improvements to the model are necessary to achieve the sensitivity and specificity required for routine clinical use, this work increases our ability to classify rare resistance mutations, thereby potentially increasing the capability of whole genome sequencing based diagnostic susceptibility testing to respond to emerging and rare resistance patterns, as well as prioritizing rare resistance mutations for *in vitro* validation. Additionally, improving the classification of susceptible *pncA* mutations will allow us to begin to disentangle the involvement of other genes in pyrazinamide resistance, including determining the effect of mutations in other pyrazinamide resistance-associated genes such as *panD* and *rpsA*.

There are two principal limitations of our approach: (1) since the training set uses a binary resistant/susceptible phenotype, the models can only predict whether a mutation confers high-level resistance (>100 µg/mL^64^) or not and (2) it can only make predictions for missense mutations in the coding sequence of *pncA*. It is known that genetic variation can lead to small changes in MIC for pyrazinamide and other first-line antitubercular compounds and that, whilst these may not change the binary phenotype, they do affect clinical outcome^65,66^. In addition, while we have shown that missense mutations represent most of the possible resistant genetic variants in *pncA*, insertions/deletions and nonsense mutations must also be considered, as they are generally associated with resistance. Likewise, promoter mutations that result in reduced transcription of *pncA* will likely also lead to resistance.

Our predictive capabilities will improve with time: the largest potential improvement is likely to come from the availability of larger datasets, preferably with pyrazinamide minimum inhibitory concentrations. Quantitative labels would help delineate mutations that result in an MIC similar to the 100 µg/mL breakpoint as one suspects that this effect is the reason why many mutations test inconsistently in the laboratory which has complicated both our training and validation. New machine learning approaches and better general-purpose predictors, especially those that aim to predict the effect of a mutation on protein stability, will no doubt come.

Even before that, predictions made by this or similar models could potentially provide clinicians with an initial estimate of pyrazinamide susceptibility after a novel mutation is observed but before traditional phenotypic testing has been completed. Given the latter can take weeks or even months, this could help guide initial therapy and further antibiotic susceptibility testing. In addition, the putative classification of additional *pncA* mutations potentially enables genetic variants conferring pyrazinamide resistance that do not involve the *pncA* gene to be discovered. The identification of pyrazinamide-susceptible mutations is also crucial, as it has been suggested that any non-synonymous mutation in *pncA* that is not cataloged as susceptible confers resistance, an incorrect assumption that would lead to overprediction of pyrazinamide resistance^67^.

The approach used here should be extensible to any pro-drug system where the enzyme is non-essential, such as delaminid, protaminid, or ethionamide, as well as to pro-drug systems in other pathogens. One promising area for future work is in the anti-tubercular bedaquiline, where resistance is caused in part by mutations in a transcriptional repressor (*Rv0678*) that cause loss of DNA binding and upregulation of efflux pumps^68,69^. Predictive methods, as shown here, will help accelerate the rate at which whole genome sequencing approaches move to the forefront of global tuberculosis control efforts.

## Author Contributions

JJC, PWF, and TMW designed experiments, CIL, DA & PWF wrote computer code used by the experiments, JJC & PWF carried out the experiments, JJC, PWF and ASW performed statistical analyses, JJC, PWF, ASW, and TEAP wrote the manuscript. PWF and DWC supervised the work. MGW, JEP and GPM contributed samples and edited the manuscript.

## Transparency Declarations

The authors have no interests to declare.

## References

1. World Health Organization. Global Tuberculosis Report. (2022).

2. Njire, M. et al. Pyrazinamide resistance in Mycobacterium tuberculosis: Review and update. Adv. Med. Sci. 61, 63–71 (2016).

3. Zhang, Y. & Yew, W. W. Mechanisms of drug resistance in Mycobacterium tuberculosis: Update 2015. Int J Tuberc Lung Dis 19, 1276–1289 (2015).

4. Zhang, Y. & Mitchison, D. The curious characteristics of pyrazinamide: A review. Int. J. Tuberc. Lung Dis. 7, 6–21 (2003).

5. Mitchison, D. A. The action of antituberculosis drugs in short-course chemotherapy. Tubercle 66, 219– 225 (1985).

6. WHO consolidated guidelines on tuberculosis: module 4: treatment: drug-susceptible tuberculosis treatment. https://www.who.int/publications-detail-redirect/9789240048126.

7. Dawson, R. et al. Efficiency and safety of the combination of moxifloxacin, pretomanid (PA-824), and pyrazinamide during the first 8 weeks of antituberculosis treatment: A phase 2b, open-label, partly randomised trial in patients with drug-susceptible or drug-resistant pul. The Lancet 385, 1738–1747 (2015).

8. Chang, K. C. et al. Pyrazinamide may improve fluoroquinolone-based treatment of multidrug-resistant tuberculosis. Antimicrob. Agents Chemother. 56, 5465–5475 (2012).

9. Zumla, A. I. et al. New antituberculosis drugs, regimens, and adjunct therapies: Needs, advances, and future prospects. Lancet Infect. Dis. 14, 327–340 (2014).

10. Nuermberger, E. et al. Powerful bactericidal and sterilizing activity of a regimen containing PA-824, moxifloxacin, and pyrazinamide in a murine model of tuberculosis. Antimicrob. Agents Chemother. 52, 1522–1524 (2008).

11. Rosenthal, I. M. et al. Daily dosing of rifapentine cures tuberculosis in three months or less in the murine model. PLoS Med. 4, 1931–1939 (2007).

12. Veziris, N. et al. A once-weekly R207910-containing regimen exceeds activity of the standard daily regimen in murine tuberculosis. Am. J. Respir. Crit. Care Med. 179, 75–79 (2009).

13. Whitfield, M. G. et al. A global perspective on pyrazinamide resistance: Systematic review and meta-analysis. PLoS ONE 10, 1–16 (2015).

14. Chang, K. C., Yew, W. W. & Zhang, Y. Pyrazinamide Susceptibility Testing in Mycobacterium tuberculosis: a Systematic Review with Meta-Analyses. Antimicrob. Agents Chemother. 55, 4499– 4505 (2011).

15. Hewlett, D., Horn, D. L. & Alfalla, C. Drug-resistant tuberculosis: inconsistent results of pyrazinamide susceptibility testing. JAMA 273, 916–7 (1995).

16. Miller, M. A., Thibert, L., Desjardins, F., Siddiqi, S. H. & Dascal, A. Testing of susceptibility of Mycobacterium tuberculosis to pyrazinamide: Comparison of Bactec method with pyrazinamidase assay. J. Clin. Microbiol. 33, 2468–2470 (1995).

17. Hoffner, S. et al. Proficiency of drug susceptibility testing of Mycobacterium tuberculosis against pyrazinamide: the Swedish experience. Int J Tuberc Lung Dis 17, 1486–1490 (2013).

18. Pandey, S., Newton, S., Upton, A., Roberts, S. & Drinkovi, D. Characterisation of pncA mutations in clinical Mycobacterium tuberculosis isolates in New Zealand. Pathology (Phila.) 41, 582–584 (2009).

19. Simons, S. O. et al. Validation of pncA gene sequencing in combination with the mycobacterial growth indicator tube method to test susceptibility of Mycobacterium tuberculosis to pyrazinamide. J. Clin. Microbiol. 50, 428–434 (2012).

20. Chedore, P., Bertucci, L., Wolfe, J., Sharma, M. & Jamieson, F. Potential for erroneous results indicating resistance when using the bactec MGIT 960 system for testing susceptibility of Mycobacterium tuberculosis to pyrazinamide. J. Clin. Microbiol. 48, 300–301 (2010).

21. World Health Organization. Catalogue of mutations in Mycobacterium tuberculosis complex and their association with drug resistance. https://www.who.int/publications/i/item/9789240028173 (2021).

22. Walker, T. M. et al. The 2021 WHO catalogue of Mycobacterium tuberculosis complex mutations associated with drug resistance: a genotypic analysis. Lancet Microbe 3, e265–e273 (2022).

23. Yüksel, P. & Tansel, Ö. Characterization of pncA mutations of pyrazinamide-resistant Mycobacterium tuberculosis in Turkey. New Microbiol. 32, 153–158 (2009).

24. Ramirez-Busby, S. M. et al. A Multinational Analysis of Mutations and Heterogeneity in PZase, RpsA, and PanD Associated with Pyrazinamide Resistance in M/XDR Mycobacterium tuberculosis. Sci. Rep. 7, 1–9 (2017).

25. Sheen, P. et al. A multiple genome analysis of Mycobacterium tuberculosis reveals specific novel genes and mutations associated with pyrazinamide resistance. BMC Genomics 18, 1–11 (2017).

26. Gopal, P. et al. Pyrazinamide resistance is caused by two distinct mechanisms: Prevention of coenzyme a depletion and loss of virulence factor synthesis. ACS Infect. Dis. 2, 616–626 (2016).

27. Zhang, Y., Zhang, J., Cui, P., Zhang, Y. & Zhang, W. Identification of Novel Efflux Proteins Rv0191, Rv3756c, Rv3008, and Rv1667c Involved in Pyrazinamide Resistance in Mycobacterium tuberculosis. Antimicrob Agent Chemo 61, e00940–17 (2017).

28. Hirano, K., Takahashi, M., Kazumi, Y., Fukasawa, Y. & Abe, C. Mutation in pncA is a major mechanism of pyrazinamide resistance in Mycobacterium tuberculosis. Tuber. Lung Dis. 78, 117–122 (1997).

29. Stoffels, K., Mathys, V., Fauville-Dufaux, M., Wintjens, R. & Bifania, P. Systematic analysis of pyrazinamide-resistant spontaneous mutants and clinical isolates of Mycobacterium tuberculosis. Antimicrob Agent Chemo 56, 5186–5193 (2012).

30. Miotto, P. et al. Mycobacterium tuberculosis pyrazinamide resistance determinants: a multicenter study. mBio 5, e01819–14 (2014).

31. Scorpio, A. & Zhang, Y. Mutations in pncA, a gene encoding pyrazinamidase/nicotinamidase, cause resistance to the antituberculous drug pyrazinamide in tubercle bacillus. Nat. Med. 2, 662–667 (1996).

32. Kim, N., Petingi, L. & Schlick, T. Network theory tools for RNA modeling. WSEAS Trans. Math. 12, 941–955 (2013).

33. Yee, M., Gopal, P. & Dick, T. Missense mutations in the unfoldase ClpC1 of the caseinolytic protease complex are associated with pyrazinamide resistance in Mycobacterium tuberculosis. Antimicrob. Agents Chemother. 61, 1–6 (2017).

34. Zhang, S. et al. Mutation in clpC1 encoding an ATP-dependent ATPase involved in protein degradation is associated with pyrazinamide resistance in Mycobacterium tuberculosis. Emerg. Microbes Infect. 6, e8–2 (2017).

35. Driesen, M. et al. Evaluation of a novel line probe assay to detect resistance to pyrazinamide, a key drug used for tuberculosis treatment. Clin. Microbiol. Infect. 24, 60–64 (2018).

36. Kalokhe, A. S. et al. Multidrug-resistant tuberculosis drug susceptibility and molecular diagnostic testing: a review of the literature. Am J Med Sci 345, 143–148 (2013).

37. Whitfield, M. G. et al. Mycobacterium tuberculosis pncA polymorphisms that do not confer pyrazinamide resistance at a breakpoint concentration of 100 micrograms per milliliter in MGIT. J. Clin. Microbiol. 53, 3633–3635 (2015).

38. Yadon, A. N. et al. A comprehensive characterization of PncA polymorphisms that confer resistance to pyrazinamide. Nat. Commun. 8, 588 (2017).

39. The CRyPTIC Consortium & 100000 Genomes Project. Prediction of Susceptibility to First-Line Tuberculosis Drugs by DNA Sequencing. New Eng J Med 379, 1403–1415 (2018).

40. Tunstall, T. et al. Combining structure and genomics to understand antimicrobial resistance. Comput. Struct. Biotechnol. J. 18, 3377–3394 (2020).

41. Davis, J. J. et al. Antimicrobial Resistance Prediction in PATRIC and RAST. Sci. Rep. 6, 1–12 (2016).

42. Deelder, W. et al. Machine learning predicts accurately mycobacterium tuberculosis drug resistance from whole genome sequencing data. Front. Genet. 10, 1–9 (2019).

43. Kouchaki, S. et al. Application of machine learning techniques to tuberculosis drug resistance analysis. Bioinformatics 35, 2276–2282 (2019).

44. Brankin, A. E. & Fowler, P. W. Predicting antibiotic resistance in complex protein targets using alchemical free energy methods. J. Comput. Chem. 43, 1771–1782 (2022).

45. Portelli, S. et al. Prediction of rifampicin resistance beyond the RRDR using structure-based machine learning approaches. Sci. Rep. 10, 1–13 (2020).

46. Karmakar, M. et al. Empirical ways to identify novel Bedaquiline resistance mutations in AtpE. PLoS ONE 14, 1–14 (2019).

47. Yang, Y. et al. DeepAMR for predicting co-occurrent resistance of Mycobacterium tuberculosis. Bioinformatics 35, 3240–3249 (2019).

48. The CRyPTIC Consortium. Quantitative drug susceptibility testing for M. tuberculosis using unassembled sequencing data and machine learning. bioRxiv (2022) doi:10.1101/2021.09.14.458035.

49. Karmakar, M., Rodrigues, C. H. M., Horan, K., Denholm, J. T. & Ascher, D. B. Structure guided prediction of Pyrazinamide resistance mutations in pncA. Sci. Rep. 10, 1–10 (2020).

50. Crook, D. W. et al. A data compendium associating the genomes of 12,289 Mycobacterium tuberculosis isolates with quantitative resistance phenotypes to 13 antibiotics. PLoS Biol. 20, e3001721 (2022).

51. Piersimoni, C. et al. Prevention of false resistance results obtained in testing the susceptibility of Mycobacterium tuberculosis to pyrazinamide with the bactec MGIT 960 system using a reduced inoculum. J. Clin. Microbiol. 51, 291–294 (2013).

52. Petrella, S. et al. Crystal structure of the pyrazinamidase of mycobacterium tuberculosis: Insights into natural and acquired resistance to pyrazinamide. PLoS ONE 6, e15785 (2011).

53. Fyfe, P. K., Rao, V. A., Zemla, A., Cameron, S. & Hunter, W. N. Specificity and mechanism of acinetobacter baumanii nicotinamidase: Implications for activation of the front-line tuberculosis drug pyrazinamide. Angew Chem Int Ed 48, 9176–9179 (2009).

54. Fowler, Philip, Lynch, Charlotte & Adlard, Dylan. https://github.com/fowler-lab/sbmlcore doi:10.5281/zenodo.8424481. (2023).

55. Rogov, S. I. & Nekrasov, A. N. A numerical measure of amino acid residues similarity based on the analysis of their surroundings in natural protein sequences. Protein Eng. 14, 459–63 (2001).

56. Blaabjerg, L. M. et al. Rapid protein stability prediction using deep learning representations. eLife 12, e82593 (2023).

57. Pires, D. E. V., Ascher, D. B. & Blundell, T. L. Structural bioinformatics mCSM: predicting the effects of mutations in proteins using graph-based signatures. 30, 335–342 (2014).

58. Cao, H., Wang, J., He, L., Qi, Y. & Zhang, J. Z. DeepDDG: Predicting the Stability Change of Protein Point Mutations Using Neural Networks. J. Chem. Inf. Model. 59, 1508–1514 (2019).

59. Stone, E. A. & Sidow, A. Physicochemical constraint violation by missense substitutions mediates impairment of protein function and disease severity. Genome Res. 15, 978–986 (2005).

60. Hecht, M., Bromberg, Y. & Rost, B. Better prediction of functional effects for sequence variants. BMC Genomics 16, S1 (2015).

61. Fowler, Philip. https://github.com/fowler-lab/predict-pyrazinamide-resistance. (2023).

62. Yoon, J. H., Nam, J. S., Kim, K. J. & Ro, Y. T. Characterization of pncA mutations in pyrazinamide-resistant Mycobacterium tuberculosis isolates from Korea and analysis of the correlation between the mutations and pyrazinamidase activity. World J. Microbiol. Biotechnol. 30, 2821–2828 (2014).

63. Fowler, P. W. et al. Robust Prediction of Resistance to Trimethoprim in Staphylococcus aureus. Cell Chem. Biol. 25, 339–349.e4 (2018).

64. World Health Organization. Technical Report on critical concentrations for drug susceptibility testing of medicines used in the treatment of drug-resistant tuberculosis. 1–128 (2018).

65. Gumbo, T. et al. The pyrazinamide susceptibility breakpoint above which combination therapy fails. J Antimicrob. Chem 69, 2420–2425 (2014).

66. Colangeli, R. et al. Bacterial Factors That Predict Relapse after Tuberculosis Therapy. New Eng J Med 379, 823–833 (2018).

67. Zignol, M. et al. Population-based resistance of Mycobacterium tuberculosis isolates to pyrazinamide and fluoroquinolones: results from a multicountry surveillance project. Lancet Infec Dis. 16, 1185–1192 (2016).

68. Milano, A. et al. Azole resistance in Mycobacterium tuberculosis is mediated by the MmpS5-MmpL5 efflux system. Tuberculosis 89, 84–90 (2009).

69. Nguyen, T. V. A., Anthony, R. M., Bañuls, A. L., Vu, D. H. & Alffenaar, J. W. C. Bedaquiline Resistance: Its Emergence, Mechanism, and Prevention. Clin. Infect. Dis. 66, 1625–1630 (2018).

