## Supplemental Information for "Prediction of pyrazinamide resistance in *Mycobacterium tuberculosis* using structure-based machine learning approaches"

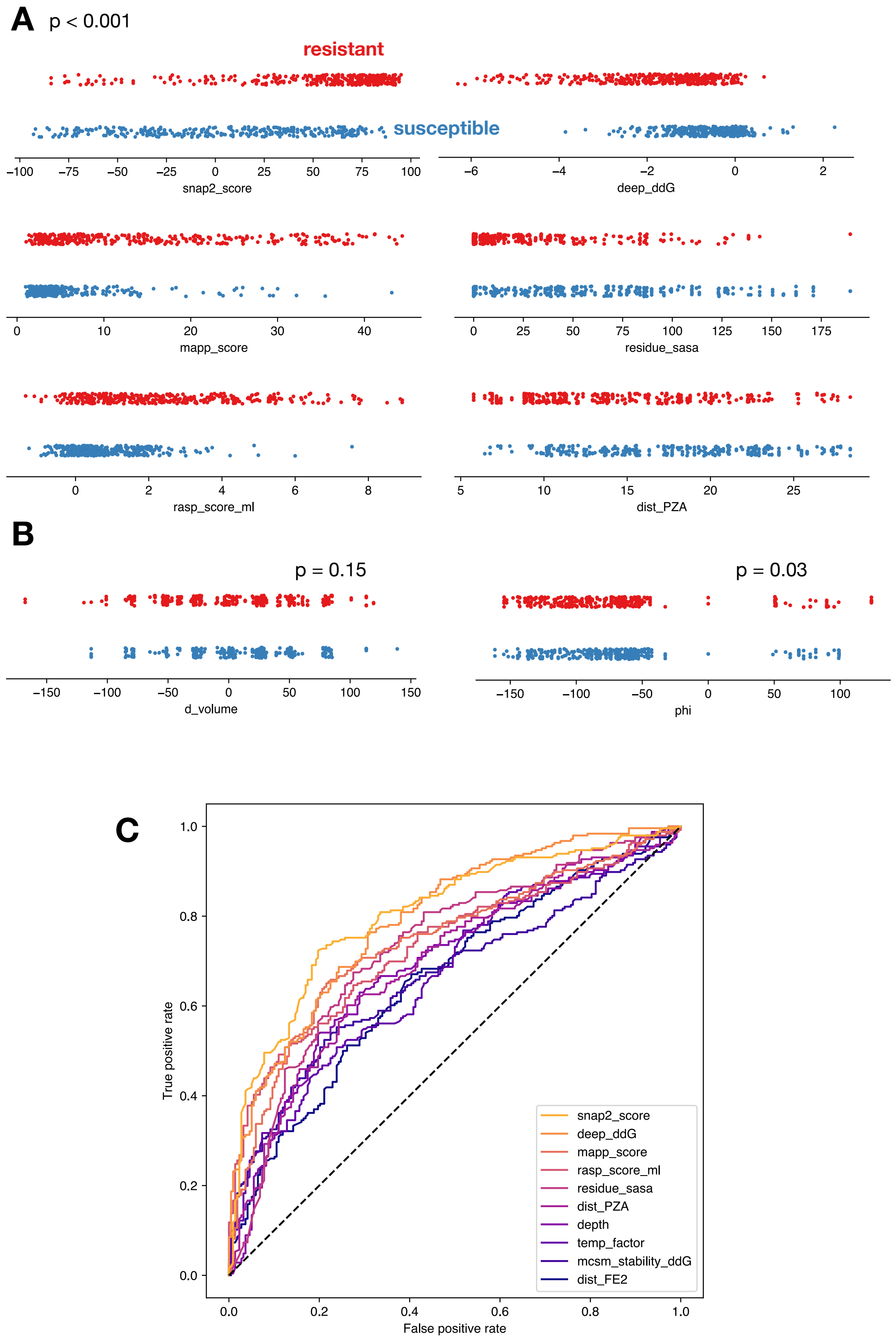


Figure S1: Related to Figure 2. Distributions of structural features vary with label. Resistant (red) and susceptible (blue) features are shown with a p-value calculated by a Mann-Whitney U test. The distributions of some features are (**A**) significantly different between resistance and susceptible mutations whilst others (**B**) are not significantly different. For clarity not all features are shown. (**C**) Receiver-operator characteristic curves for all features with an area under the curve $> 0.55$ after training a logistic regression model on each feature in turn.

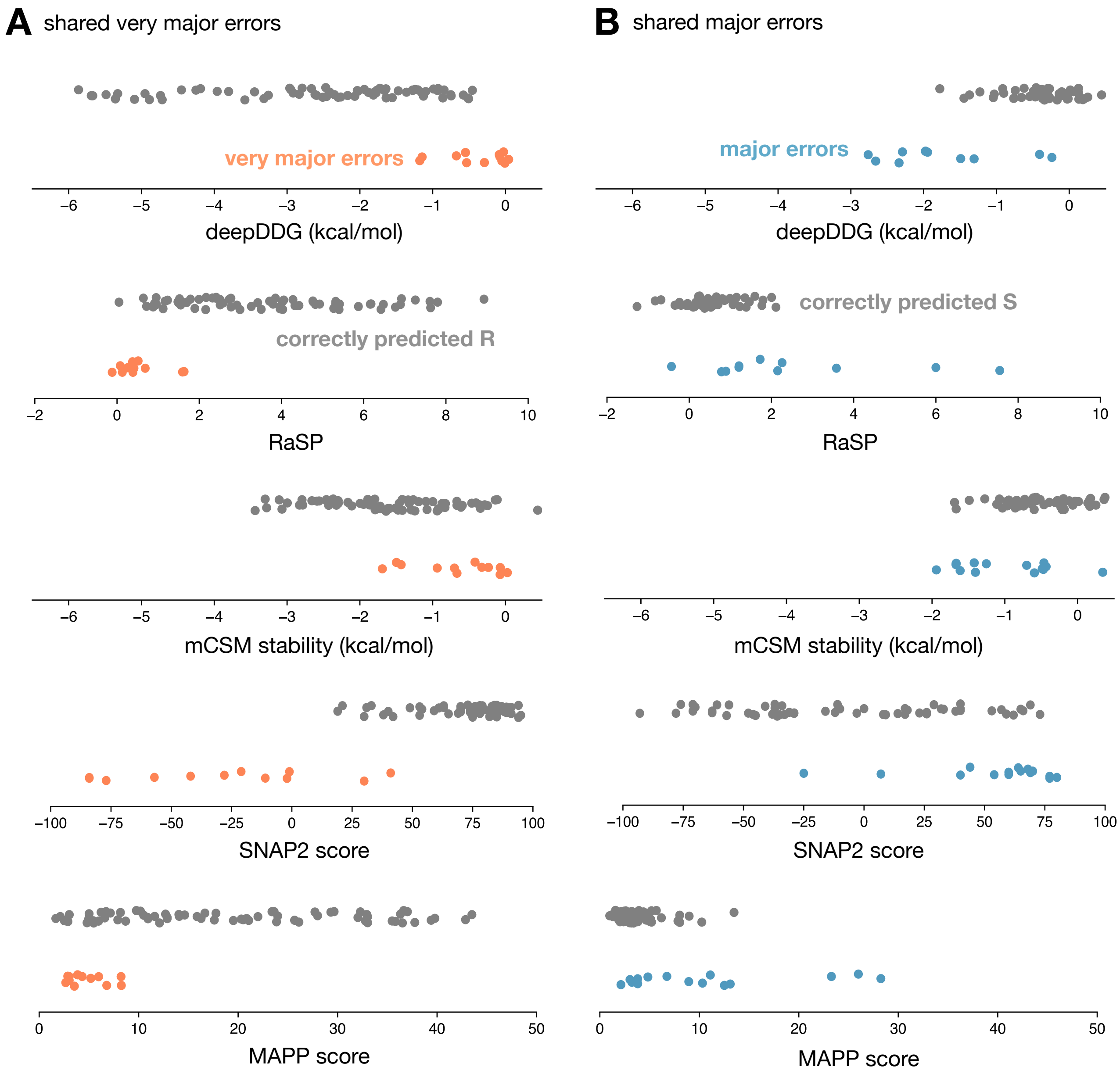


Figure S2: Related to Figure 4. The prediction of the (**A**) very major errors (orange) and (**B**) major errors (blue) shared between the three machine learning models is driven by a number of features.

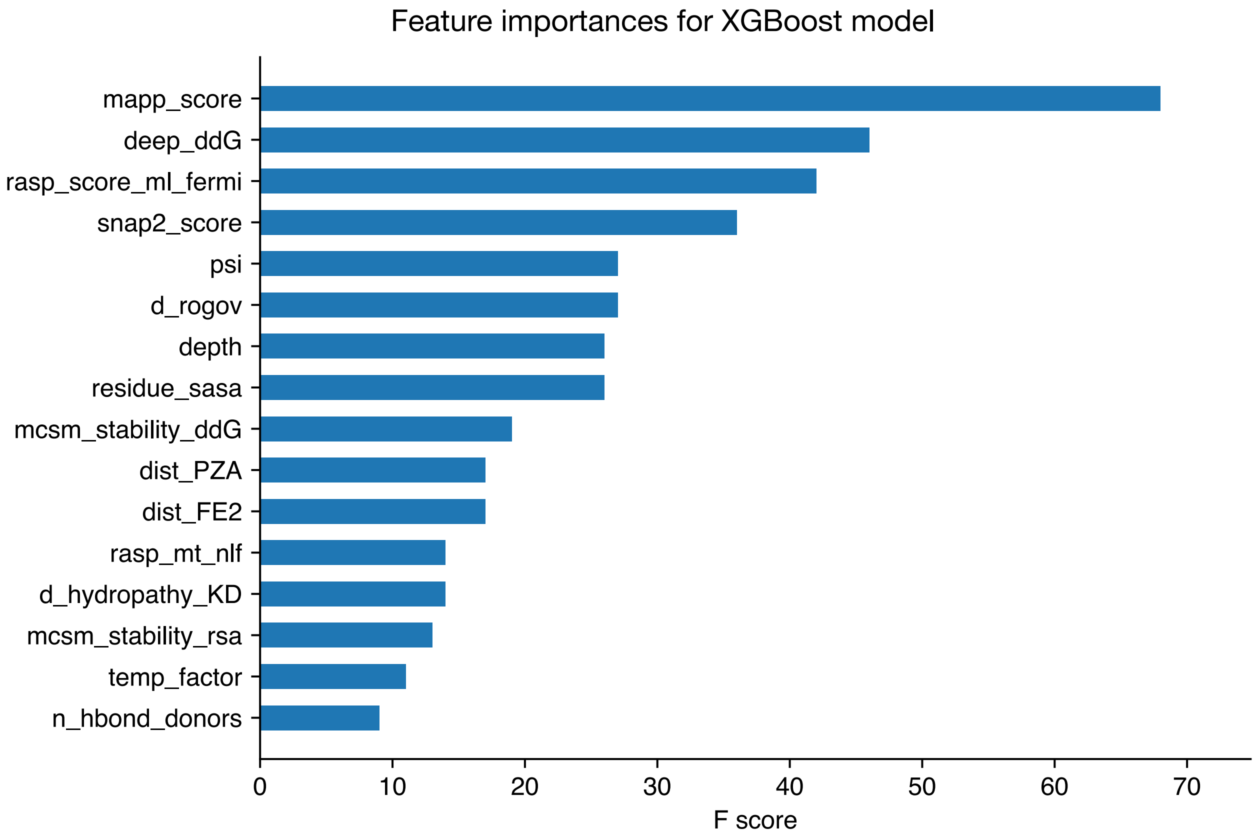


Figure S3: The gradient-boosted decision tree model makes use of a wide range of features.

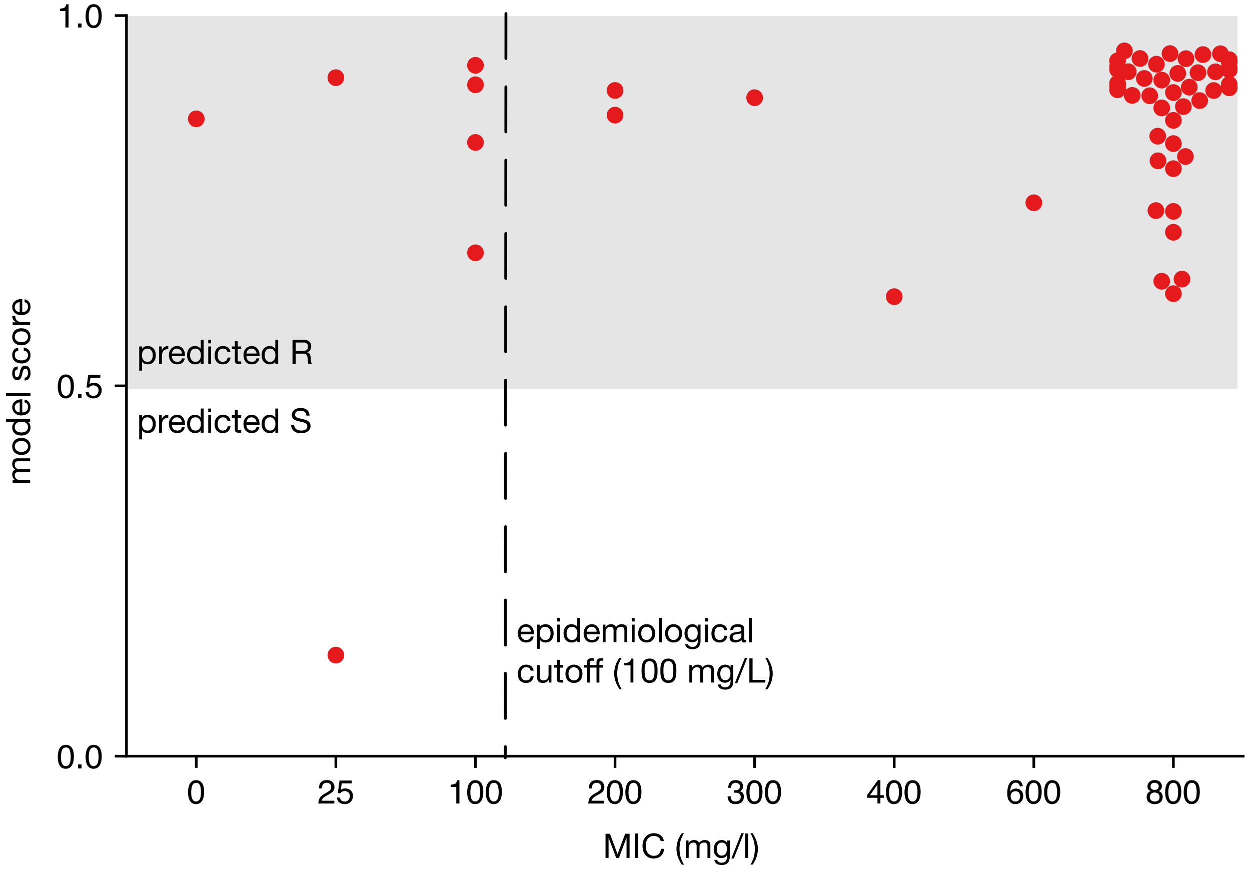


Figure S4: The gradient-boosted decision tree model correctly predicts 51 out of 57 mutations, however the dataset contains mainly resistant-conferring mutations. The epidemiological cutoff is the highest minimum inhibitory concentration observed within a phenotypically wild-type population; here that is 100 mg/L and is represented by the dashed line, however this has been shifted to make it clear that the four samples with an MIC of 100 mg/L are experimentally susceptible.
